# Neural signatures of automatic letter–speech sound integration in literate adults

**DOI:** 10.1101/2025.06.09.658589

**Authors:** Weiyong Xu, Xueqiao Li, Orsolya Kolozsvari, Aino Sorsa, Miriam Nokia, Jarmo Hämäläinen

**Affiliations:** Department of Psychology, University of Jyväskylä, Jyväskylä, Finland; Jyväskylä Centre for Interdisciplinary Brain Research, University of Jyväskylä, Jyväskylä, Finland

**Keywords:** Letter–speech sound integration, Automaticity, Learning, Magnetoencephalography (MEG), Multivariate pattern analysis (MVPA)

## Abstract

Automaticity in decoding print is crucial for fluent reading. This process relies on associative memories between letters and speech sounds (LSS) that are overlearned through years of reading practice. While previous neuroimaging studies have identified neural correlates of LSS integration across different stages of reading development, the specific neural signatures underlying automatic LSS integration remain unclear. In the present study, we aimed to isolate neural components specifically associated with automatic LSS integration in literate adults. To this end, we developed an artificial script training paradigm in which adult native Finnish speakers were taught to associate unfamiliar foreign letters with familiar Finnish speech sounds. Using magnetoencephalography (MEG), we directly compared the audiovisual processing of newly learned and overlearned LSS associations within the same task, one day after training. Event-related fields (ERFs) and multivariate decoding revealed largely shared neural circuits of audiovisual integration for both types of LSS associations, as evidenced by multisensory interaction and congruency effects. Interestingly, the processing of congruent overlearned audiovisual associations uniquely recruited brain activity in the left parietal cortex during the 235-475 ms time window. Furthermore, temporal generalization analysis of the congruency effects revealed that while both newly learned and overlearned audiovisual associations engaged common neural mechanisms, the newly learned associations were processed systematically more slowly by a few hundred milliseconds. Our study identified the spatiotemporal neural signatures underlying automatized LSS processing, offering insights into neural markers that may help identify levels of reading proficiency.

## 1. Introduction

Learning the correspondence between letters and sounds is a fundamental step in learning to read in alphabetic languages (Blomert & Froyen, 2010; Treiman et al., 1998). Although most children acquire letter knowledge within the first year of schooling (D. J. W. Froyen et al., 2009; Sigmundsson et al., 2020), achieving automaticity in the neural processing of letter–speech sound (LSS) associations takes years of reading practice (Blomert & Froyen, 2010; D. J. W. Froyen et al., 2009; Joo et al., 2019; Xu et al., 2018). The transition from slow, laborious decoding to rapid, effortless LSS transformation is crucial for fluent reading (Joo et al., 2019; Romanovska & Bonte, 2021; van Atteveldt & Ansari, 2014) and failure to automate the grapheme-to-phoneme mapping process has been linked to reading difficulties in alphabetic languages (Blau et al., 2010; Romanovska et al., 2021; Wang et al., 2020; Žarić et al., 2014).

Extensive research has demonstrated that reading practice reorganizes brain circuits for visual processing (Dehaene et al., 2010; Romanovska & Bonte, 2021). The left ventral occipitotemporal (vOT) cortex plays a crucial role in the development of reading skills, showing functional specialization for print processing and language integration (Lerma-Usabiaga et al., 2018; Price & Devlin, 2011). According to the neuronal recycling hypothesis, the left vOT becomes specialized for letter and word recognition by repurposing a portion of the ventral visual pathway originally dedicated to face and object recognition (Dehaene & Cohen, 2011). The biased connectivity hypothesis further suggests that the consistent involvement of the left vOT region in reading results from pre-existing connections between this subregion and areas involved in spoken-language processing (Bouhali et al., 2014; Hannagan et al., 2021; Mahon & Caramazza, 2011; Saygin et al., 2016). Recent evidence suggests that reading experience drives this reorganization within the left vOT, contributing to its role in linking visual print with spoken language (Debska et al., 2024; Taylor et al., 2019). Similarly, LSS integration has been conceptualized as an assimilation and accommodation process within the superior temporal cortices (STC), where neural substrates initially used for speech-sound and object-sound integration are adapted for LSS integration (D. Gao et al., 2024). Neuroimaging studies consistently identify the left STC as a central region for LSS integration (Blau et al., 2010; C. Gao et al., 2023; Holloway et al., 2015; Van Atteveldt et al., 2004; van Atteveldt et al., 2009). Several other brain regions, including the vOT (I Karipidis et al., 2017; Karipidis et al., 2021; Pleisch et al., 2019), inferior frontal gyrus (IFG) (Hashimoto & Sakai, 2004; Karipidis et al., 2021; Li et al., 2020), and parietal areas (Raij et al., 2000; Xia et al., 2022; Xu et al., 2018) are likewise considered parts of the core brain network responsible for LSS integration.

In addition to the emergence of specialized cortical networks for LSS processing, studies using high-temporal-resolution techniques, such as electroencephalography (EEG) and MEG, have identified specific time windows for LSS integration in the brain (Caffarra et al., 2021; D. Froyen et al., 2010; Jost et al., 2014; Raij et al., 2000; Xu et al., 2018, 2019). One line of research on the temporal dynamics in LSS integration comes from studies using cross-modal mismatch negativity (MMN) paradigms. One early MMN study found that the simultaneous presentation of letters enhances the MMN response to deviant speech sounds in literate adults, suggesting early and automatic integration (D. Froyen et al., 2008). In contrast, beginner readers (one year of reading experience) showed differences only in later time-windows (∼650 ms) and no MMN for LSS integration (D. J. W. Froyen et al., 2009). Furthermore, congruent LSS, where the visual letter matches the corresponding speech sound (for example, seeing “b” and hearing /b/), affected both early MMN and later ERPs (P300), suggesting multiple stages of processing (Andres et al., 2011). Additionally, when looking at dyslexic adults, audiovisual integration was deficient and sluggish compared to fluent readers (Andres et al., 2011; Blomert & Froyen, 2010; Mittag et al., 2013). However, using MMN to study audiovisual integration is limited, as it primarily reflects the automatic deviance detection mechanisms and does not directly measure the integration of audiovisual stimuli (O’Reilly & O’Reilly, 2021). Further, MMN cannot capture the full range of processes involved in audiovisual integration, particularly the dynamics that depend on top-down cognitive control or attentional mechanisms (Paraskevopoulos et al., 2012). Therefore, although automatic LSS integration has been inferred using the MMN paradigm, more direct evidence from integration tasks examining the temporal dynamics of automatic LSS processing in the brain is still lacking.

LSS integration has been assessed using various analytical approaches across studies (C. Gao et al., 2023; Richlan, 2019) including congruency effects (audiovisual congruent vs. incongruent), interaction effects (audiovisual vs. the sum of auditory and visual responses) and conjunction-based analyses. However, there is considerable variability in the criteria used to detect LSS integration, often leading to inconsistent findings regarding the associated brain regions and time windows (C. Gao et al., 2023). For example, interaction effects based on the additive model are straightforward for electrophysiological studies (Senkowski et al., 2011; Stein & Stanford, 2008), but specific analytical contrasts employed in fMRI studies differ substantially (Beck et al., 2024; Karipidis et al., 2021; Van Atteveldt et al., 2004; Xia et al., 2022). Moreover, LSS integration has been shown to be sensitive to various factors, including imaging modality, experimental design, language transparency, and developmental stage (Beck et al., 2024; Holloway et al., 2015; Karipidis et al., 2021; Pleisch et al., 2019; Wang et al., 2020; Xia et al., 2022). Consequently, there does not appear to be a consistent or stable neural signature of automatic LSS integration. Supporting this, congruency effect—the most simple and commonly used measure—could be observed even after short training sessions with artificial LSS associations in both adults (J. A. Hämäläinen et al., 2019; Xu et al., 2020) and in children (I Karipidis et al., 2017), suggesting that it may not necessarily reflect automatic LSS integration.

A key strategy for understanding automatic LSS integration involves examining how reading experience transforms its underlying neural network beyond the initial learning phase. While early learning of arbitrary LSS associations likely relies on general multisensory memory systems (Murray & Shams, 2023; Vogt, 2023), overlearned processing is thought to engage specialized cortical circuits and become automatic through years of reading practice (Karipidis et al., 2021; Romanovska & Bonte, 2021). However, automatic LSS representations typically emerge alongside brain maturation and cognitive development, making it difficult to disentangle the specific effects of reading experience. Artificial LSS training offers a controlled framework to simulate early learning stages and explore the experience-dependent plasticity of LSS integration (Aravena et al., 2017; Elbro et al., 2012; Folia et al., 2010; J. A. Hämäläinen et al., 2019). In our previous study (Xu et al., 2020), we trained literate adults to associate unfamiliar letters with Finnish phonemes and observed dynamic, unstable neural representations of newly learned LSS associations across two days of training.

In this study, we aimed to investigate the neural network involved in automatic LSS integration by examining the differences in MEG brain activities between newly learned and overlearned LSS associations in literate adults. First, we aimed to replicate and validate the overlearned audiovisual processing in transparent languages, as reported by earlier studies (Karipidis et al., 2021; Raij et al., 2000; Van Atteveldt et al., 2004; Xu et al., 2018). We then directly compared the processing of newly learned (Xu et al., 2020) and overlearned audiovisual associations in the brain using MEG data from the same experiment, thereby controlling for factors that typically influence audiovisual integration. By applying source localization of event-related fields and multivariate pattern analysis, we aimed to identify spatiotemporal neural signatures that reflect the automaticity of LSS integration. Additionally, we expected to observe differences in brain activations to unimodally presented overlearned and newly learned letters (Fernández-López et al., 2021; Joo et al., 2019). We hypothesized that literate adults are able to reuse neural circuits involved in processing overlearned LSS associations when learning new ones (Dehaene et al., 2010; Martin et al., 2019). Therefore, we expected similar brain regions to be recruited for audiovisual integration in newly learned transparent LSS associations. However, we hypothesized that overlearned audiovisual associations would be processed more automatically and rapidly, possibly supported by specialized cortical neural circuits for storing and processing overlearned letter to sound mappings. The audiovisual interaction process, as indexed by the additive model, was expected to be similar for both types of associations, given that it involves more general purpose cross-modal interactions (Calvert & Calvert, 2001; Xu et al., 2019).

## 2. Methods

### 2.1 Participants

In total, 36 participants were recruited through email-lists and posters. The majority of them were young university students and staff. Data from four participants was not collected because of cancellation. The remaining 32 participants (22 females, 2 left handed, 2 ambidextrous, mean age 24.22 years, SD 3.43 years, range 19-36 years) were included in this study. Participants were prescreened according to the following exclusion criteria: head injuries, neurological diseases, attention deficit hyperactivity disorder (ADHD), medication usage that could affect the central nervous system, language delays, or other kind of language-related impairment. All of them had normal hearing and normal or corrected-to-normal vision based on self-report. Ethical approval was obtained from the Ethics Committee of the University of Jyväskylä, and the study was conducted in accordance with the Declaration of Helsinki. Written informed consent was obtained from all the participants prior to their participation in the study. The participants were given gift cards or movie tickets (comparable value of 30 euros) as compensation for their time spent in the MEG experiments and cognitive tests sessions.

### 2.2 Stimuli and task

Auditory stimuli consisted of 12 Finnish phonemes ([a], [ä], [e], [t], [s], [k], [o], [ö], [i], [p], [v], [d]; mean duration: 473 ms; SD:103 ms). Visual stimuli consisted of 12 Georgian letters (□, ჵ, ჹ, უ, დ, ჱ, ც, ჴ, ნ, ფ, ღ, წ) and 12 Finnish letters (a, ä, e, t, s, k, o, ö, i, p, v, d). The Georgian letters were divided into two sets with 6 stimuli in each set. Each participant was trained during two days to associate one set (Learnable set) of the Georgian letters with 6 Finnish phonemes, while they did not learn the corresponding sounds for the other set (Control set). The two letter sets (Learnable and Control, Figure 1B) were counterbalanced between the participants. See our previous study (Xu et al., 2020) for a detailed description of the training procedure.

**Figure 1.**
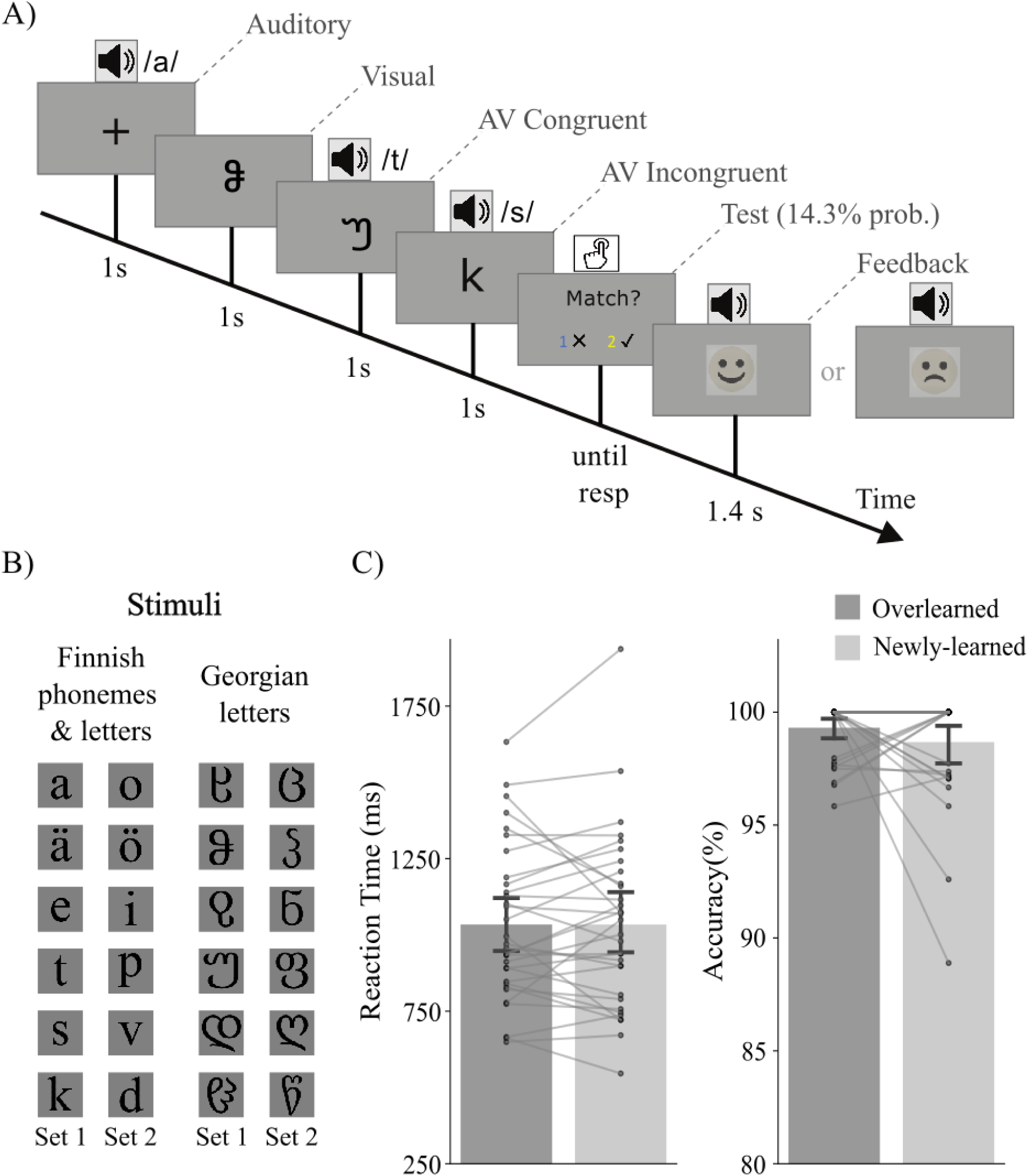
Experimental design, stimuli, and behavioral performance in the audiovisual task. A) Schematic illustration of the experimental design. The four trial types (A, V, AVC, and AVI) were presented in random order. B) Stimuli used in the experiment, including Finnish letters and phonemes (overlearned) and Georgian letters (newly learned). Two sets of stimuli (Set 1 and Set 2) were created and used counterbalanced across participants. C) Behavioral performance in the audiovisual task. There were no significant differences between the overlearned and newly learned conditions for reaction time and accuracy.

After the above-mentioned two training sessions on two consecutive days, they participated in an audiovisual integration task (Figure 1A) after the training session on the second day. In the audiovisual integration task both newly learned (Georgian letter-sound pairs from the Learnable set) and overlearned (Finnish letter-sound pairs, which shared the same phonemes as the newly learned Georgian letter-sound pairs) audiovisual stimuli were presented randomly. The Finnish letter-sound associations were overlearned since the participants were native Finnish literate adults. There were also auditory only and visual only trials in which only a unimodal stimulus was presented. Each trial started with a 1-second fixation cross followed by a 1-second auditory only, visual only, or simultaneously presented audiovisual stimulus. To make the participants focus on the stimuli, there were test trials which occurred at a 14.3% probability following an audiovisual stimulus. They were instructed to press the correct button on a response pad to indicate whether the auditory and visual stimuli were congruent or incongruent. Congruency here refers to whether the auditory stimulus corresponded with the visual stimulus or not. The two buttons corresponding to correct and incorrect answers were randomly generated for each test trial. The test trial stayed on the screen until the participant made a response. After the response, they were given audiovisual feedback (1400 ms) on the accuracy of their response.

A short practice session was presented before the formal experiment. Two Georgian and two Finnish LSS pairs were used for practice (in total 16 trials). These LSS pairs were also used for practice in the previous two training sessions. The experiment consisted of 6 blocks. In each block, there were two repetitions of visual (V) and audiovisual congruent (AVC) and audiovisual incongruent (AVI) stimuli for the newly learned and overlearned conditions. The auditory only (A) condition was also repeated twice in each block. Therefore, there were 72 trials for each condition across the whole experiment. After each block, accuracy of their responses in the current block and overall accuracy during the test were presented to the participants to keep them motivated in the task. They were also instructed to take a short break after completing each block.

### 2.3 Data acquisition and pre-processing

The magnetoencephalography (MEG) data was recorded in a magnetically shielded room at the Centre for Interdisciplinary Brain Research, University of Jyväskylä, using the 306-channel (102 magnetometers and 204 gradiometers) Elekta Neuromag® TRIUX™ system. To monitor the participant’s head position within the MEG helmet, five digitized head position indicator (HPI) coils were attached to the head: three on the forehead and one behind each ear. Before the MEG experiment, three anatomical landmarks (nasion, left and right preauricular points), the position of the five HPI coils, and the overall head shape of the participants (>100 points evenly distributed over the scalp) were digitized with the Polhemus Isotrak digital tracker system (Polhemus, Colchester, VT, United States). Electrooculogram (EOG) was measured with two electrodes placed diagonally around the eyes, one slightly below the left eye and the other slightly above the right eye. A ground electrode was attached to the collarbone. The sampling rate was set to 1000 Hz, and a 0.1–330 Hz online band-pass filter was applied during the MEG data acquisition. The MEG machine was in a 68° upright gantry position, and the participants were comfortably seated in a chair during data collection.

Raw MEG data were first preprocessed with Maxfilter (version 3.0.17) with the movement compensated temporal signal-space separation (tSSS) method (Taulu & Simola, 2006) to suppress the noise interference that originated outside of the head and to compensate for the signal distortion related to head movement. Bad (noisy or flat) MEG channels were checked manually and marked before Maxfilter; the signals in the bad channels were then reconstructed by Maxfilter.

After Maxfilter, MEG Data were processed in MNE Python (Gramfort et al., 2013) (version: 1.6.1). First, noisy segments of MEG data were annotated manually and were excluded from further analysis. A band pass filter of 0.1-40 Hz (zero-phase FIR filter design using the “hamming” window method) was applied to the continuous MEG data. Fast independent component analysis (FastICA) was then used to remove cardiac and eye movement related artifacts (Hyvärinen, 1999). Data was then segmented into epochs from −200 ms to 1000 ms relative to the stimulus onset. A peak-to-peak amplitude (grad = 1500e-13 T/m, mag = 5e-12 T) rejection threshold was used to remove any bad epochs and then the epochs were visually inspected in case of some remaining artifacts. Baseline correction was applied to each channel and epoch by subtracting the mean signal of the baseline period (−200 ms to 0 ms relative to the stimulus onset) from the entire epoch.

### 2.4 Data analysis

#### 2.4.1. Linear regression-based ERF analysis

To facilitate comparison with previous studies on LSS integration, we first analyzed the ERF responses in our dataset. We employed linear regression analysis (implemented using the “linear_regression_raw” function in MNE Python) to examine the evoked responses of audiovisual interaction [AV − (A+V)] (Besle et al., 2004, 2008) and congruency (AVC-AVI) effects (Raij et al., 2000; Xu et al., 2020). Specifically, the audiovisual congruent response was constructed as the sum of unisensory auditory (A), unisensory visual (V), audiovisual interaction effect (Interaction), and congruency effect (Xu et al., 2020):

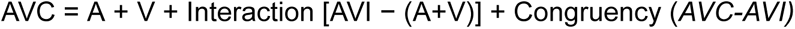

Linear regression-based evoked responses were calculated for newly learned (NL) and overlearned (OL) conditions separately: OL_V, NL_V, OL_interaction, NL_interaction, OL_congruency, and NL_congruency.

#### 2.4.2 Source reconstruction

Since individual MRIs were not collected in this study, the Freesurfer (RRID: SCR_001847, Martinos Center for Biomedical Imaging, Charlestown, MA, United States) average brain (FsAverage) was used as template for MEG source analysis. The template brain was scaled uniformly and coregistered to the digitized head points of the participants using an automated approach described in (Houck & Claus, 2020). A surface-based source space was used with an ‘ico5’ spacing parameter (10242 sources per hemisphere). Forward solution was calculated by using a single-compartment boundary-element model (BEM). Depth-weighted (p = 0.8) minimum-norm estimates (wMNE) (M. S. Hämäläinen & Ilmoniemi, 1994; Lin et al., 2006) were calculated with loose constraint source orientation. Dynamic statistical parametric maps (dSPM) (Dale et al., 2000) were used for noise normalization and the source activities normal to the cortical surface were used for further analysis.

#### 2.4.3. Multivariate pattern analysis

##### Temporal decoding

A logistic regression classifier with L2 regularization (C=1.0) was implemented using scikit-learn’s ‘liblinear’ solver (max iterations = 1000). Binary classifiers were trained on epochs from two conditions (e.g., congruent vs. incongruent). Subsampling was employed to equalize the number of trials across the two conditions, ensuring balanced class distributions. Decoders were trained at each time point of the epoch, utilizing the amplitude of 306 MEG channels as features. These features were standardized by removing the mean and scaling to unit variance across epochs using the sklearn.preprocessing.StandardScaler function. Classifier performance was evaluated using the area under the receiver operating characteristic curve (ROC-AUC) score with 5-fold cross-validation. The weight vectors from the classifiers were transformed into activation patterns, which are more neurophysiologically interpretable (Haufe et al., 2014). The activation patterns were visualized as a topographical map to assess the importance of each channel in terms of its contribution to the decoding performance.

##### Temporal generalization

Temporal generalization (TG) (King & Dehaene, 2014) was used to further characterize the temporal organization of the different audiovisual processing stages. Basically, the trained classifiers at each time point are tested on its ability to generalize to all time points in the same condition (within-condition TG), or from one experimental condition to another (cross-condition TG). This procedure produces a two-dimensional temporal generalization matrix, offering a more nuanced perspective than simple latency analyses by revealing how neural representations emerge, evolve, or stabilize over time.

#### 2.4.4. Statistical analysis

Paired t-tests on the behavioral data were conducted using the stats module from the SciPy package (Virtanen et al., 2020). Permutation t-tests with spatio-temporal clustering were used for the statistical analysis of dSPM source analysis of the ERF data (Maris & Oostenveld, 2007). To avoid the issue of specifying a free yet somewhat arbitrary threshold for the initial clustering, the threshold-free cluster enhancement method (TFCE: h_power=2.00, e_power=0.50, start=0, step=0.2) was utilized (Smith & Nichols, 2009). The time window for the comparison between the brain activation to newly learned and overlearned visual stimuli and audiovisual interaction effects was set to 0 ms to 1000 ms post stimulus onset. The time window for the congruency effects was set to 200 ms to 1000 ms post stimulus onset based on previous studies (Caffarra et al., 2021; Raij et al., 2000; Xu et al., 2019). For statistical analysis of the decoding results, similar temporal clustering permutation tests were used on the decoding scores against chance level. The number of permutations was set to 1024, and the statistical alpha level was set at 0.05 for all tests.

## 3. Results

### 3.1 Behavioral performance for the test trials

There was no significant difference in reaction time between overlearned LSS pairs (1015 ± 256 ms) and newly learned LSS pairs (1021 ± 288 ms; t = 0.20, p = 0.84). Accuracy also did not differ between overlearned LSS pairs (0.992 ± 0.013) and newly learned LSS pairs (0.986 ± 0.026; t = 1.09, p = 0.29). See Figure 1C for a summary plot of the behavioral performance.

### 3.2 Unimodal differences between newly learned and overlearned letters

#### 3.2.1 Enhanced ERF source activity for newly learned letters

Source-level ERF activity showed significant differences between newly learned and overlearned letters in the time window of 110–995 ms (p < 0.05, TFCE-corrected). These effects were primarily observed in the left and right superior temporal cortex (STC), ventral occipitotemporal cortex (voT), medial temporal lobe (MTL), and left parietal cortex (PC), with stronger source activity for newly learned letters compared to overlearned letters (Figure 2A).

**Figure 2.**
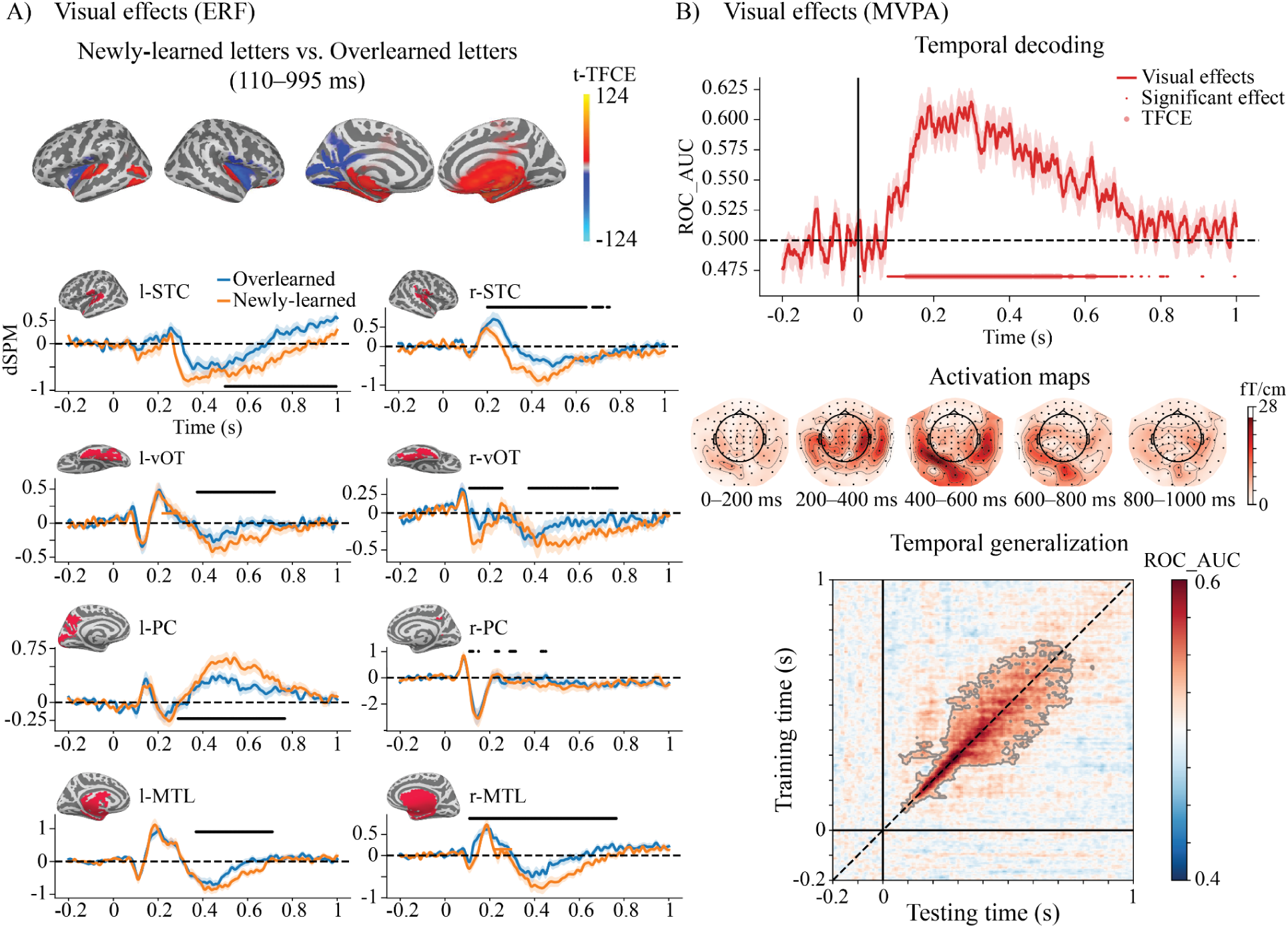
Differences in visual processing of newly learned and overlearned letters. A) ERF source results showing the difference between newly learned and overlearned letters. The upper panels display the TFCE-t maps of the significant effects averaged over the significant time windows (p < 0.05, TFCE-corrected). The lower panels visualize the significant differences with dSPM waveforms extracted from eight brain regions showing prominent differences. B) MVPA results for the newly learned and overlearned letters. The upper panel shows the AUC-ROC scores for temporal decoding. The middle panel visualizes the activation maps of the decoders trained over time. The lower panel shows the temporal generalization of the decoding, with significant generalization highlighted in darker shades (p < 0.05, TFCE-corrected).

#### 3.2.2 Presence of two distinct visual processes in temporal generalization

Comparable differences in visual responses to newly learned and overlearned letters were also observed using temporal decoding (Figure 2B, top). The AUC-ROC scores were significantly higher than chance level in the time window of 128–624 ms (p < 0.05, TFCE-corrected). The activation maps of the significant time points were visualized as topographic maps (Figure 2B, middle). The temporal generalization matrix (Figure 2B, bottom) showed that classification accuracy was highest along the diagonal when the training time points were close to the testing time points, particularly in the early time window around 100–300 ms. After 300 ms, generalization appeared to broaden, extending over a time window of approximately 200 ms. These TG results suggest the presence of at least two distinct visual processes: an early, sequential process characterized by rapidly changing transient representations and a more sustained process at later time points after 300 ms.

### 3.3 Audiovisual interaction effects

Significant audiovisual interaction effects [AVI − (A+V)] (p < 0.05, TFCE-corrected) were observed for both newly learned (30–995 ms) and overlearned LSS associations (65–995 ms) in distributed brain regions (see Figure 3A). No differences in the interaction effects were found between the two types of LSS associations.

**Figure 3.**
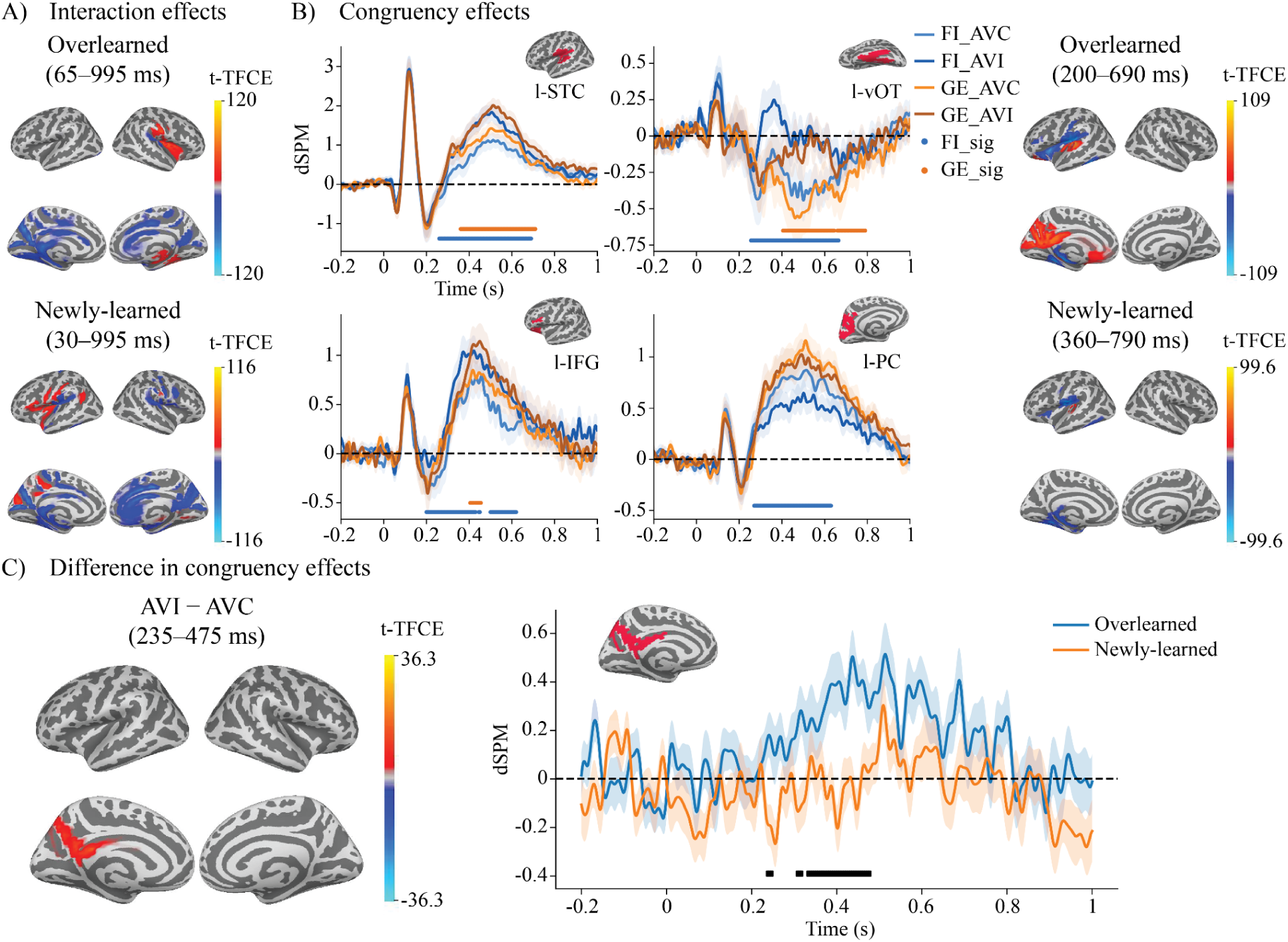
ERF results of audiovisual interaction and congruency effects for newly learned and overlearned LSS associations. A) Interaction effects were observed for both newly learned and overlearned LSS associations in distributed brain regions (p < 0.05, TFCE-corrected). No difference in interaction effects was found between the two types of LSS associations. B) Significant congruency effects were found for both newly learned and overlearned LSS associations (p < 0.05, TFCE-corrected), primarily in the left STC, vOT, PC, and IFG. The right panel visualizes the significant differences with dSPM waveforms extracted from four brain regions showing prominent differences. C) A unique LSS congruency process was found near the left parietal cortex (PC), specific to overlearned LSS processing, during the time window of 235–475 ms. The right panel visualizes the significant differences with dSPM waveforms extracted from the left PC.

### 3.4 Audiovisual congruency effects

#### 3.4.1 ERF source activity reveals distinct and common congruency effects between overlearned and newly learned LSS associations

We first investigated the audiovisual congruency effect for overlearned LSS and newly learned LSS separately. Significant ERF congruency effects (p < 0.05, TFCE-corrected) were found for both newly learned (360–790 ms) and overlearned (200–690 ms) LSS associations (Figure 3B). Both congruency effects were primarily localized in the left STC, vOT, IFG, and parietal cortex. We then compared the congruency effects between overleaned and newly learned conditions to examine any learning-related differences. Interestingly, when comparing the congruency effect between overlearned and newly learned LSS associations, a unique LSS congruency effect (p < 0.05, TFCE-corrected) was observed near the left parietal region, specific to overlearned LSS processing, during the relatively early time window of 235–475 ms (235–245 ms, 305–315 ms, and 330–475 ms) (Figure 3C).

#### 3.4.2 Decoding analysis suggests newly learned LSS associations are processed slower and longer compared to overlearned LSS associations

Both newly learned and overlearned congruency effects exhibited above-chance decoding scores (overlearned: 373–660 ms; newly learned: 543–997 ms, Figure 4A). Temporal decoding activation maps (Figure 4B) revealed that significant decoding scores were driven by brain activity in the left temporal channels. The temporal generalization profiles showed distinct patterns for newly learned and overlearned associations: overlearned congruency information was processed more rapidly and in a structured manner, whereas newly learned congruency processing was slower, involving a distinct, later sustained process (Figure 4C, top row). Notably, cross-condition TG analysis revealed that the two audiovisual processes were decodable from each other, with the processing of newly learned LSS consistently appearing slower, as indicated by the asymmetric generalization pattern along the diagonal (Figure 4C, bottom row).

**Figure 4.**
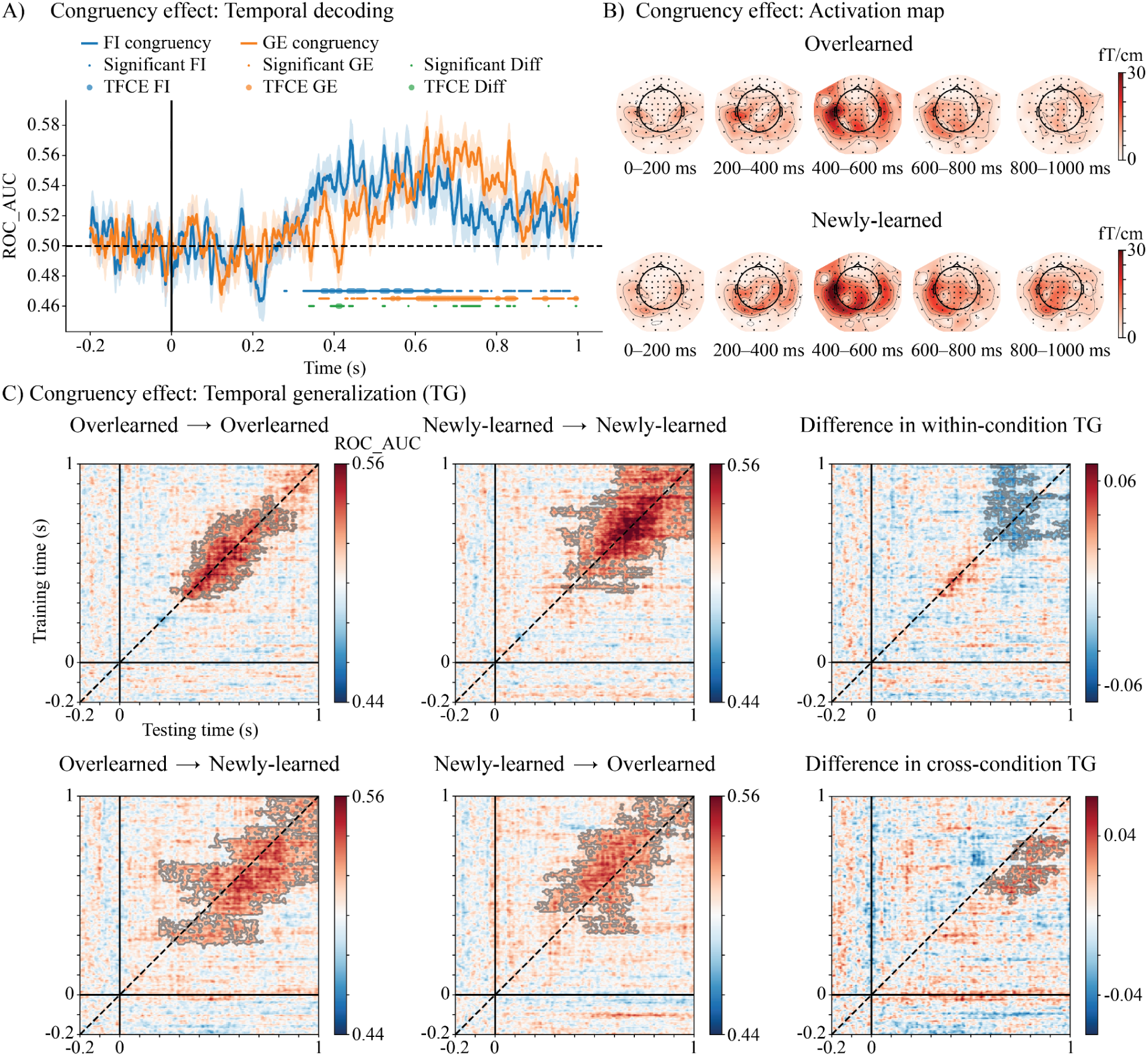
MVPA results for the congruency effects of newly learned and overlearned audiovisual associations. A) Temporal decoding of congruency effects for newly learned and overlearned associations. B) Activation maps of the decoders trained over time. C) Temporal generalization (TG) of congruency effects within conditions (overlearned → overlearned, newly learned → newly learned) and between conditions (overlearned → newly learned, newly learned → overlearned). The third column displays the difference in TG scores for within-condition and between-condition comparisons. Significant generalization is highlighted in darker shades (p < 0.05, TFCE-corrected)

## 4. Discussion

In this study, we systematically investigated the neural signature of the automaticity of LSS integration using both ERF analyses and MVPA. These two complementary approaches revealed, for the first time, consistent evidence that long-term reading experience fine-tunes the brain network contributing towards audiovisual integration to support the automatic processing of overlearned LSS associations in literate adults. Although behavioral performance in the cross-modal matching task did not significantly differ between newly learned and overlearned associations after two days of training, neural responses showed clear distinctions in both visual and audiovisual processing. Specifically, newly learned and overlearned letters evoked distinct neural activations extending beyond visual areas into auditory and memory-related regions. While the core network supporting audiovisual integration was largely shared across learning levels, congruent overlearned LSS associations uniquely engaged the left parietal cortex at 235–475 ms post-stimulus. Temporal generalization analysis further revealed a shared neural representation of congruency across different learning levels, but with a notable difference in processing speed: Brain responses to newly learned LSS associations were consistently delayed by several hundred milliseconds compared to overlearned ones.

We observed distinct spatiotemporal cortical activation patterns for newly learned and overlearned letters when presented unimodally. Overall, newly learned letters elicited larger ERF responses than overlearned letters in the bilateral ventral occipito-temporal cortex (vOT), superior temporal cortex (STC), parietal regions, and the medial temporal lobe (MTL). Temporal generalization analysis further revealed an early stage of sequential processing with rapidly evolving neural representations between 100–300 ms, followed by a later stage (300–800 ms) characterized by more stable and sustained activity patterns (Thesen et al., 2012). The increased activation in the vOT aligns with the interactive account of the vOT function (Price & Devlin, 2011), which suggests that heightened activation during early learning reflects greater prediction error, gradually diminishing as experience accumulates. This pattern is consistent with electrophysiological findings showing an inverted U-shaped developmental trajectory of the N1/N170 response to print, with selectivity peaking during the early stages of learning and declining in skilled readers (Fraga-González et al., 2021; Maurer et al., 2006). Furthermore, it is also in line with previous studies (Brem et al., 2018; Pleisch et al., 2019; Xu et al., 2020) demonstrating that training with artificial character–speech sound associations leads to increased activation for the trained characters within the vOT. Our results extend previous findings by showing that experience-dependent differences in letter processing manifest across a distributed brain network. The observed differences likely reflect the less consolidated audiovisual memory representations of newly learned letters, which consequently rely more heavily on the MTL (Staresina & Wimber, 2019) for retrieval of their phonemic representations and demand greater processing resources and broader cortical network engagement.

The audiovisual interaction, referring to the brain’s ability to combine auditory and visual information, was observed under both overlearned and newly learned conditions, with no significant difference between them. The associated neural activation patterns were widely distributed across the brain, a finding consistent with our previous study (Xu et al., 2018).These AV interaction effects are considered to reflect more basic cross-modal integration processes, which may be less influenced by reading experience. Furthermore, the relatively early onset latency of these effects aligns with prior studies documenting early cross-modal integration mechanisms (Molholm et al., 2002; Raij et al., 2024). It is also possible that the observed interactions reflect additional cognitive processes, such as attention, engaged during the active task in our study. The distributed brain activation patterns seem to suggest a mixed contribution from both basic sensory integration and higher-level cognitive processing.

Most interestingly, a significant audiovisual congruency effect emerged in the left hemisphere in both newly learned and overlearned conditions. The overlearned congruency effects in the STC, vOT, parietal cortex and IFG are largely consistent with previous studies on LSS integration (C. Gao et al., 2023; Karipidis et al., 2021; Raij et al., 2000; van Atteveldt & Ansari, 2014; Wang et al., 2020; Xia et al., 2022; Xu et al., 2019). A novel insight from our study is the functional specialization we identified within this network. While the newly learned congruency effect was found in a largely similar brain network as the overlearned congruency, a direct contrast between the congruency effects in the two learning conditions revealed a difference specifically within a left parietal region near the precuneus. This finding is intriguing in light of previous research indicating that posterior neocortical areas, such as the precuneus and angular gyrus, are not only involved in perceptual information processing but are also associated with long-term memory storage after memory consolidation (Krenz et al., 2023). This is consistent with the observations in the present study that the congruency effect was absent in the precuneus for newly learned associations whereas it was stronger for the overlearned congruent LSS associations than for the incongruent during the 235–475 ms time window. The posterior parietal cortex (PPC) has been shown to play a significant role in the development and performance of arbitrary visuomotor associations, particularly as these mappings become overlearned (Grol et al., 2006; Madhavan et al., 2019). Research indicates that while the PPC is not essential for the initial learning of these associations, its involvement becomes more pronounced with extensive training, which can enhance the automaticity of learned mappings (Grol et al., 2006, 2009).

In addition to identifying a specialized cortical site in the left parietal cortex for processing overlearned LSS associations, the ERF results also suggest that the significant temporal window for congruency effects seems to occur several hundred milliseconds earlier for overlearned compared to newly learned LSS associations, particularly in the left STC and vOT (Figure 3B). Converging and more direct evidence from decoding and temporal generalization analyses further supports a difference in processing speed between these two types of LSS associations. The within-condition temporal generalization revealed distinct patterns: overlearned LSS integration was processed more efficiently, as indicated by a narrower off-diagonal generalization pattern, whereas newly learned LSS integration was processed more slowly and involved a distinct sustained activity in the later time window. This difference in processing speed is also reflected in the cross-condition temporal generalization analysis, which revealed an asymmetric generalization pattern along the diagonal of the temporal generalization matrix (King & Dehaene, 2014).These multiple analyses consistently indicate a robust difference in the temporal dynamics of newly learned versus overlearned LSS integration. Furthermore, cross-condition decoding analyses provided compelling evidence for a shared neural representation underlying the LSS integration process, as multivariate classifiers trained to distinguish congruent from incongruent stimuli successfully generalized between newly learned and overlearned conditions.Taken together, these findings suggest that the congruency effect is not directly related to modality-specific sensory input but instead reflects an abstract representation of audiovisual congruency, with a distinct temporal dynamic between newly learned and overlearned LSS.

In general, the findings from this study revealed distinct spatiotemporal brain processes related to automaticity in audiovisual integration, which may stem from the involvement of different neural circuits for processing newly learned LSS associations compared to overlearned LSS associations. According to the systems consolidation theory of memory (Klinzing et al., 2019; Squire et al., 2015), newly learned associations initially rely on the medial temporal lobe (MTL) and gradually become integrated into the neocortex. It is therefore likely that learned letter–sound associations are first supported by domain-general associative learning networks, with hippocampal involvement during the early, effortful mapping of orthographic and phonological representations (Servant et al., 2018). With continued exposure and practice, these associations are gradually consolidated into specialized neocortical circuits, enabling automatic and efficient processing (Gilboa & Moscovitch, 2021; Sekeres et al., 2018). For learning the novel LSS associations in literate adults, it seems that the brain utilizes a large part of the general audiovisual integration network, yet the level of automaticity remains lower compared to overlearned LSS associations. The faster integration speed of overlearned LSS associations may be due to the involvement of additional cortical regions in the left parietal cortex for storing and processing these associations.

In this study, we examined the learning of new LSS associations in literate adults who have already developed an automatic LSS integration network through decades of reading experience. A parallel can be drawn from research on musicians, who, through extensive training in note–sound associations, exhibit distinct neural and behavioral advantages that extend even to novel audiovisual learning. This suggests that long-term experience and overlearning may facilitate more efficient processing and the ability to generalize learned associations across different domains (Paraskevopoulos et al., 2012, 2014). Building on these findings, it would be intriguing to extend this research to children at varying reading skill levels, using artificial LSS learning paradigms (I Karipidis et al., 2017) to investigate the spatiotemporal development of automaticity in congruency effects, as identified in our study. Furthermore, examining dyslexic readers with similar MVPA and time-sensitive neuroimaging methods could offer valuable insights into whether they exhibit specific impairments in the automatic integration of LSS (D. J. W. Froyen et al., 2009).

In conclusion, these findings highlight how reading experience continues to shape neural circuits for audiovisual integration, enabling faster and more efficient processing of audiovisual information. Our results suggest that in literate adults, learning new LSS associations largely engages the same reading network used for processing overlearned associations, but with greater effort and at a slower speed. Crucially, our study reveals that automaticity is not merely about strengthening existing pathways; it also involves the specialization of distinct neural circuits, such as the left parietal cortex, which form dedicated “shortcuts” for storing and processing overlearned associations. This provides a new perspective on the neural basis of skilled reading and offers potential neural markers for tracking the transition from effortful learning to automatic proficiency.

## Data and code availability

The dataset cannot be publicly shared due to legal restrictions but can be obtained from the corresponding author upon request, subject to a formal data-sharing agreement. Under the GDPR, MEG and EEG data are classified as pseudonymous personal data and, therefore, cannot be openly distributed. In line with the 2017 guidelines from the Finnish Data Protection Ombudsman, only fully anonymized data may be publicly available. Analysis scripts are available at the GitHub repository (https://github.com/weiyongxu/AVTest.git).

## Author contributions

Weiyong Xu: Conceptualization, Methodology, Software, Formal analysis, Investigation, Data Curation, Writing - Original Draft, Writing - Review & Editing, Visualization

Aino Sorsa: Investigation, Writing - Review & Editing

Orsolya Kolozsvari: Conceptualization, Writing - Review & Editing

Xueqiao Li: Visualization, Writing - Review & Editing

Miriam Nokia: Supervision, Writing - Review & Editing

Jarmo Hämäläinen: Conceptualization, Supervision, Writing - Review & Editing

## Acknowledgments

We would like to thank Suvi Karjalainen and Ainomaija Laitinen for their help in data collection.

## Reference

Andres, A. J. D., Oram Cardy, J. E., & Joanisse, M. F. (2011). Congruency of auditory sounds and visual letters modulates mismatch negativity and P300 event-related potentials. International Journal of Psychophysiology: Official Journal of the International Organization of Psychophysiology, 79(2), 137–146.

Aravena, S., Tijms, J., Snellings, P., & Molen, M. W. van der. (2017). Predicting Individual Differences in Reading and Spelling Skill With Artificial Script–Based Letter–Speech Sound Training. Journal of Learning Disabilities, 0022219417715407.

Beck, J., Chyl, K., Dębska, A., Łuniewska, M., van Atteveldt, N., & Jednoróg, K. (2024). Letter-speech sound integration in typical reading development during the first years of formal education. Child Development, 95(4), e236–e252.

Besle, J., Fischer, C., Bidet-Caulet, A., Lecaignard, F., Bertrand, O., & Giard, M.-H. (2008). Visual Activation and Audiovisual Interactions in the Auditory Cortex during Speech Perception: Intracranial Recordings in Humans. Journal of Neuroscience, 28(52), 14301–14310.

Besle, J., Fort, A., Delpuech, C., & Giard, M.-H. (2004). Bimodal speech: early suppressive visual effects in human auditory cortex. The European Journal of Neuroscience, 20(8), 2225–2234.

Blau, V., Reithler, J., van Atteveldt, N., Seitz, J., Gerretsen, P., Goebel, R., & Blomert, L. (2010). Deviant processing of letters and speech sounds as proximate cause of reading failure: A functional magnetic resonance imaging study of dyslexic children. Brain: A Journal of Neurology, 133(Pt 3), 868–879.

Blomert, L., & Froyen, D. (2010). Multi-sensory learning and learning to read. International Journal of Psychophysiology: Official Journal of the International Organization of Psychophysiology, 77(3), 195–204.

Bouhali, F., Thiebaut de Schotten, M., Pinel, P., Poupon, C., Mangin, J.-F., Dehaene, S., & Cohen, L. (2014). Anatomical connections of the visual word form area. The Journal of Neuroscience: The Official Journal of the Society for Neuroscience, 34(46), 15402–15414.

Brem, S., Hunkeler, E., Mächler, M., Kronschnabel, J., Karipidis, I. I., Pleisch, G., & Brandeis, D. (2018). Increasing expertise to a novel script modulates the visual N1 ERP in healthy adults. International Journal of Behavioral Development, 42(3), 333–341.

Caffarra, S., Lizarazu, M., Molinaro, N., & Carreiras, M. (2021). Reading-related brain changes in audiovisual processing: cross-sectional and longitudinal MEG evidence. The Journal of Neuroscience: The Official Journal of the Society for Neuroscience. 10.1523/JNEUROSCI.3021-20.2021

Calvert, G. a., & Calvert, G. a. (2001). Crossmodal processing in the human brain: insights from functional neuroimaging studies. Cerebral Cortex, 11, 1110–1123.

Dale, A. M., Liu, A. K., Fischl, B. R., Buckner, R. L., Belliveau, J. W., Lewine, J. D., & Halgren, E. (2000). Dynamic statistical parametric mapping: combining fMRI and MEG for high-resolution imaging of cortical activity. Neuron, 26(1), 55–67.

Debska, A., Wang, S., Jednorog, K., & Pattamadilok, C. (2024). Reading acquisition drives linguistic cross-modal convergence in the left ventral occipitotemporal cortex. In bioRxiv (p. 2024.11.26.625405). 10.1101/2024.11.26.625405

Dehaene, S., & Cohen, L. (2011). The unique role of the visual word form area in reading. Trends in Cognitive Sciences, 15(6), 254–262.

Dehaene, S., Pegado, F., Braga, L. W., Ventura, P., Nunes Filho, G., Jobert, A., Dehaene-Lambertz, G., Kolinsky, R., Morais, J., & Cohen, L. (2010). How learning to read changes the cortical networks for vision and language. Science, 330(6009), 1359–1364.

Elbro, C., Daugaard, H. T., & Gellert, A. S. (2012). Dyslexia in a second language?-a dynamic test of reading acquisition may provide a fair answer. Annals of Dyslexia, 62(3), 172–185.

Fernández-López, M., Marcet, A., & Perea, M. (2021). Does orthographic processing emerge rapidly after learning a new script? British Journal of Psychology, 112(1), 52–91.

Folia, V., Uddén, J., De Vries, M., Forkstam, C., & Petersson, K. M. (2010). Artificial language learning in adults and children. Language Learning, 60(s2), 188–220.

Fraga-González, G., Pleisch, G., Di Pietro, S. V., Neuenschwander, J., Walitza, S., Brandeis, D., Karipidis, I. I., & Brem, S. (2021). The rise and fall of rapid occipito-temporal sensitivity to letters: Transient specialization through elementary school. Developmental Cognitive Neuroscience, 49, 100958.

Froyen, D. J. W., Bonte, M. L., van Atteveldt, N., & Blomert, L. (2009). The long road to automation: neurocognitive development of letter-speech sound processing. Journal of Cognitive Neuroscience, 21(3), 567–580.

Froyen, D., van Atteveldt, N., & Blomert, L. (2010). Exploring the Role of Low Level Visual Processing in Letter-Speech Sound Integration: A Visual MMN Study. Frontiers in Integrative Neuroscience, 4, 9.

Froyen, D., Van Atteveldt, N., Bonte, M., & Blomert, L. (2008). Cross-modal enhancement of the MMN to speech-sounds indicates early and automatic integration of letters and speech-sounds. Neuroscience Letters, 430(1), 23–28.

Gao, C., Green, J. J., Yang, X., Oh, S., Kim, J., & Shinkareva, S. V. (2023). Audiovisual integration in the human brain: a coordinate-based meta-analysis. Cerebral Cortex (New York, N.Y.: 1991), 33(9), 5574–5584.

Gao, D., Liang, X., Ting, Q., Nichols, E. S., Bai, Z., Xu, C., Cai, M., & Liu, L. (2024). A meta-analysis of letter-sound integration: Assimilation and accommodation in the superior temporal gyrus. Human Brain Mapping, 45(15), e26713.

Gilboa, A., & Moscovitch, M. (2021). No consolidation without representation: Correspondence between neural and psychological representations in recent and remote memory. Neuron, 109(14), 2239–2255.

Gramfort, A., Luessi, M., Larson, E., Engemann, D. A., Strohmeier, D., Brodbeck, C., Goj, R., Jas, M., Brooks, T., Parkkonen, L., & Hämäläinen, M. (2013). MEG and EEG data analysis with MNE-Python. Frontiers in Neuroscience, 7, 267.

Grol, M. J., de Lange, F. P., Verstraten, F. A. J., Passingham, R. E., & Toni, I. (2006). Cerebral changes during performance of overlearned arbitrary visuomotor associations. The Journal of Neuroscience: The Official Journal of the Society for Neuroscience, 26(1), 117–125.

Grol, M. J., Toni, I., Lock, M., & Verstraten, F. A. J. (2009). Spatial representation of overlearned arbitrary visuomotor associations. Experimental Brain Research, 192(4), 751–759.

Hämäläinen, J. A., Parviainen, T., Hsu, Y.-F., & Salmelin, R. (2019). Dynamics of brain activation during learning of syllable-symbol paired associations. Neuropsychologia, 129, 93–103.

Hämäläinen, M. S., & Ilmoniemi, R. J. (1994). Interpreting magnetic fields of the brain: minimum norm estimates. Medical & Biological Engineering & Computing, 32(1), 35–42.

Hannagan, T., Agrawal, A., Cohen, L., & Dehaene, S. (2021). Emergence of a compositional neural code for written words: Recycling of a convolutional neural network for reading. Proceedings of the National Academy of Sciences of the United States of America, 118(46). 10.1073/pnas.2104779118

Hashimoto, R., & Sakai, K. L. (2004). Learning letters in adulthood: Direct visualization of cortical plasticity for forming a new link between orthography and phonology. Neuron, 42(2), 311–322.

Haufe, S., Meinecke, F., Görgen, K., Dähne, S., Haynes, J.-D., Blankertz, B., & Bießmann, F. (2014). On the interpretation of weight vectors of linear models in multivariate neuroimaging. NeuroImage, 87, 96–110.

Holloway, I. D., Van Atteveldt, N., Blomert, L., & Ansari, D. (2015). Orthographic dependency in the neural correlates of reading: Evidence from audiovisual integration in english readers. Cerebral Cortex, 25(6), 1544–1553.

Houck, J. M., & Claus, E. D. (2020). A comparison of automated and manual co-registration for magnetoencephalography. PloS One, 15(4), e0232100.

Hyvärinen, A. (1999). Fast and robust fixed-point algorithms for independent component analysis. IEEE Transactions on Neural Networks / a Publication of the IEEE Neural Networks Council, 10(3), 626–634.

I Karipidis, I., Pleisch, G., Röthlisberger, M., Hofstetter, C., Dornbierer, D., Stämpfli, P., & Brem, S. (2017). Neural initialization of audiovisual integration in prereaders at varying risk for developmental dyslexia. Human Brain Mapping, 38(2), 1038–1055.

Joo, S. J., Tavabi, K., Caffarra, S., & Yeatman, J. D. (2019). Automaticity in the reading circuitry. Brain and Language, 214, 829937.

Jost, L. B., Eberhard-Moscicka, A. K., Frisch, C., Dellwo, V., & Maurer, U. (2014). Integration of Spoken and Written Words in Beginning Readers: A Topographic ERP Study. Brain Topography, 27(6), 786–800.

Karipidis, I. I., Pleisch, G., Di Pietro, S. V., Fraga-González, G., & Brem, S. (2021). Developmental trajectories of letter and speech sound integration during reading acquisition. Frontiers in Psychology, 12, 750491.

King, J.-R., & Dehaene, S. (2014). Characterizing the dynamics of mental representations: the temporal generalization method. Trends in Cognitive Sciences, 18(4), 203–210.

Klinzing, J. G., Niethard, N., & Born, J. (2019). Mechanisms of systems memory consolidation during sleep. Nature Neuroscience, 22(10), 1598–1610.

Krenz, V., Alink, A., Sommer, T., Roozendaal, B., & Schwabe, L. (2023). Time-dependent memory transformation in hippocampus and neocortex is semantic in nature. Nature Communications, 14(1), 1–17.

Lerma-Usabiaga, G., Carreiras, M., & Paz-Alonso, P. M. (2018). Converging evidence for functional and structural segregation within the left ventral occipitotemporal cortex in reading. Proceedings of the National Academy of Sciences of the United States of America, 115(42), E9981–E9990.

Lin, F.-H., Witzel, T., Ahlfors, S. P., Stufflebeam, S. M., Belliveau, J. W., & Hämäläinen, M. S. (2006). Assessing and improving the spatial accuracy in MEG source localization by depth-weighted minimum-norm estimates. NeuroImage, 31(1), 160–171.

Li, Y., Seger, C., Chen, Q., & Mo, L. (2020). Left inferior frontal gyrus integrates multisensory information in category learning. Cerebral Cortex (New York, N.Y.: 1991), 30(8), 4410–4423.

Madhavan, R., Bansal, A. K., Madsen, J. R., Golby, A. J., Tierney, T. S., Eskandar, E. N., Anderson, W. S., & Kreiman, G. (2019). Neural interactions underlying visuomotor associations in the human brain. Cerebral Cortex (New York, N.Y.: 1991), 29(11), 4551–4567.

Mahon, B. Z., & Caramazza, A. (2011). What drives the organization of object knowledge in the brain? Trends in Cognitive Sciences, 15(3), 97–103.

Maris, E., & Oostenveld, R. (2007). Nonparametric statistical testing of EEG- and MEG-data. Journal of Neuroscience Methods, 164(1), 177–190.

Martin, L., Hirshorn, E. A., Durisko, C., Moore, M. W., Schwartz, R., Zheng, Y., & Fiez, J. A. (2019). Do adults acquire a second orthography using their native reading network? Journal of Neurolinguistics, 50, 120–135.

Maurer, U., Brem, S., Kranz, F., Bucher, K., Benz, R., Halder, P., Steinhausen, H. C., & Brandeis, D. (2006). Coarse neural tuning for print peaks when children learn to read. NeuroImage, 33(2), 749–758.

Mittag, M., Thesleff, P., Laasonen, M., & Kujala, T. (2013). The neurophysiological basis of the integration of written and heard syllables in dyslexic adults. Clinical Neurophysiology: Official Journal of the International Federation of Clinical Neurophysiology, 124(2), 315–326.

Molholm, S., Ritter, W., Murray, M. M., Javitt, D. C., Schroeder, C. E., & Foxe, J. J. (2002). Multisensory auditory–visual interactions during early sensory processing in humans: a high-density electrical mapping study. Cognitive Brain Research, 14(1), 115–128.

Murray, C. A., & Shams, L. (2023). Crossmodal interactions in human learning and memory. Frontiers in Human Neuroscience, 17, 1181760.

O’Reilly, J. A., & O’Reilly, A. (2021). A critical review of the deviance detection theory of mismatch negativity. NeuroSci, 2(2), 151–165.

Paraskevopoulos, E., Kuchenbuch, A., Herholz, S. C., Foroglou, N., Bamidis, P., & Pantev, C. (2014). Tones and numbers: A combined EEG-MEG study on the effects of musical expertise in magnitude comparisons of audiovisual stimuli. Human Brain Mapping, 35(11), 5389–5400.

Paraskevopoulos, E., Kuchenbuch, A., Herholz, S. C., & Pantev, C. (2012). Musical expertise induces audiovisual integration of abstract congruency rules. Journal of Neuroscience, 32(50), 18196–18203.

Pleisch, G., Karipidis, I. I., Brauchli, C., Röthlisberger, M., Hofstetter, C., Stämpfli, P., Walitza, S., & Brem, S. (2019). Emerging neural specialization of the ventral occipitotemporal cortex to characters through phonological association learning in preschool children. NeuroImage, 189, 813–831.

Price, C. J., & Devlin, J. T. (2011). The interactive account of ventral occipitotemporal contributions to reading. Trends in Cognitive Sciences, 15(6), 246–253.

Raij, T., Lin, F.-H., Letham, B., Lankinen, K., Nayak, T., Witzel, T., Hämäläinen, M., & Ahveninen, J. (2024). Onset timing of letter processing in auditory and visual sensory cortices. Frontiers in Integrative Neuroscience, 18, 1427149.

Raij, T., Uutela, K., & Hari, R. (2000). Audiovisual integration of letters in the human brain. Neuron, 28(2), 617–625.

Richlan, F. (2019). The Functional Neuroanatomy of Letter-Speech Sound Integration and Its Relation to Brain Abnormalities in Developmental Dyslexia. Frontiers in Human Neuroscience, 13. 10.3389/fnhum.2019.00021

Romanovska, L., & Bonte, M. (2021). How learning to read changes the listening brain. Frontiers in Psychology, 12, 726882.

Romanovska, L., Janssen, R., & Bonte, M. (2021). Cortical responses to letters and ambiguous speech vary with reading skills in dyslexic and typically reading children. NeuroImage. Clinical, 30(102588), 102588.

Saygin, Z. M., Osher, D. E., Norton, E. S., Youssoufian, D. A., Beach, S. D., Feather, J., Gaab, N., Gabrieli, J. D. E., & Kanwisher, N. (2016). Connectivity precedes function in the development of the visual word form area. Nature Neuroscience, 19(9), 1250–1255.

Sekeres, M. J., Winocur, G., & Moscovitch, M. (2018). The hippocampus and related neocortical structures in memory transformation. Neuroscience Letters, 680, 39–53.

Senkowski, D., Saint-Amour, D., Höfle, M., & Foxe, J. J. (2011). Multisensory interactions in early evoked brain activity follow the principle of inverse effectiveness. NeuroImage, 56(4), 2200–2208.

Servant, M., Cassey, P., Woodman, G. F., & Logan, G. D. (2018). Neural bases of automaticity. Journal of Experimental Psychology. Learning, Memory, and Cognition, 44(3), 440–464.

Sigmundsson, H., Haga, M., Ofteland, G. S., & Solstad, T. (2020). Breaking the reading code: Letter knowledge when children break the reading code the first year in school. New Ideas in Psychology, 57(100756), 100756.

Smith, S. M., & Nichols, T. E. (2009). Threshold-free cluster enhancement: Addressing problems of smoothing, threshold dependence and localisation in cluster inference. NeuroImage, 44(1), 83–98.

Squire, L. R., Genzel, L., Wixted, J. T., & Morris, R. G. (2015). Memory consolidation. Cold Spring Harbor Perspectives in Biology, 7(8), a021766.

Staresina, B. P., & Wimber, M. (2019). A neural chronometry of memory recall. Trends in Cognitive Sciences, 23(12), 1071–1085.

Stein, B. E., & Stanford, T. R. (2008). Multisensory integration: current issues from the perspective of the single neuron. Nature Reviews. Neuroscience, 9(4), 255–266.

Taulu, S., & Simola, J. (2006). Spatiotemporal signal space separation method for rejecting nearby interference in MEG measurements. Physics in Medicine and Biology, 51(7), 1759–1768.

Taylor, J. S. H., Davis, M. H., & Rastle, K. (2019). Mapping visual symbols onto spoken language along the ventral visual stream. Proceedings of the National Academy of Sciences, 201818575.

Thesen, T., McDonald, C. R., Carlson, C., Doyle, W., Cash, S., Sherfey, J., Felsovalyi, O., Girard, H., Barr, W., Devinsky, O., Kuzniecky, R., & Halgren, E. (2012). Sequential then interactive processing of letters and words in the left fusiform gyrus. Nature Communications, 3(1), 1284.

Treiman, R., Tincoff, R., Rodriguez, K., Mouzaki, A., & Francis, D. J. (1998). The foundations of literacy: learning the sounds of letters. Child Development, 69(6), 1524–1540.

van Atteveldt, N., & Ansari, D. (2014). How symbols transform brain function: A review in memory of Leo Blomert. Trends in Neuroscience and Education, 3(2), 44–49.

Van Atteveldt, N., Formisano, E., Goebel, R., & Blomert, L. (2004). Integration of letters and speech sounds in the human brain. Neuron, 43(2), 271–282.

van Atteveldt, N., Roebroeck, A., & Goebel, R. (2009). Interaction of speech and script in human auditory cortex: Insights from neuro-imaging and effective connectivity. Hearing Research, 258(1-2), 152–164.

Virtanen, P., Gommers, R., Oliphant, T. E., Haberland, M., Reddy, T., Cournapeau, D., Burovski, E., Peterson, P., Weckesser, W., Bright, J., van der Walt, S. J., Brett, M., Wilson, J., Millman, K. J., Mayorov, N., Nelson, A. R. J., Jones, E., Kern, R., Larson, E., … SciPy 1.0 Contributors. (2020). SciPy 1.0: fundamental algorithms for scientific computing in Python. Nature Methods, 17(3), 261–272.

Vogt, K. (2023). Neuroscience: Merging multisensory memories. Current Biology: CB, 33(15), R817–R819.

Wang, F., Karipidis, I. I., Pleisch, G., Fraga-González, G., & Brem, S. (2020). Development of Print-Speech Integration in the Brain of Beginning Readers With Varying Reading Skills. Frontiers in Human Neuroscience, 14, 289.

Xia, Z., Yang, T., Cui, X., Hoeft, F., Liu, H., Zhang, X., Shu, H., & Liu, X. (2022). Neurofunctional basis underlying audiovisual integration of print and speech sound in Chinese children. The European Journal of Neuroscience, 55(3), 806–826.

Xu, W., Kolozsvari, O. B., Monto, S. P., & Hämäläinen, J. A. (2018). Brain responses to letters and speech sounds and their correlations with cognitive skills related to reading in children. Frontiers in Human Neuroscience, 12, 304.

Xu, W., Kolozsvari, O. B., Oostenveld, R., & Hämäläinen, J. A. (2020). Rapid changes in brain activity during learning of grapheme-phoneme associations in adults. NeuroImage, 220, 117058.

Xu, W., Kolozsvári, O. B., Oostenveld, R., Leppänen, P. H. T., & Hämäläinen, J. A. (2019). Audiovisual processing of Chinese characters elicits suppression and congruency effects in MEG. Frontiers in Human Neuroscience, 13, 18.

Žarić, G., Fraga González, G., Tijms, J., van der Molen, M. W., Blomert, L., & Bonte, M. (2014). Reduced neural integration of letters and speech sounds in dyslexic children scales with individual differences in reading fluency. PloS One, 9(10), e110337.

